# Feature selectivity of corticocortical feedback along the primate dorsal visual pathway

**DOI:** 10.1101/2024.02.21.581426

**Authors:** Yavar Korkian, Nardin Nakhla, Christopher C. Pack

## Abstract

Anatomical studies have revealed a prominent role for feedback projections in the primate visual cortex. Theoretical models suggest that these projections support important brain functions, like attention, prediction, and learning. However, these models make different predictions about the relationship between feedback connectivity and neuronal stimulus selectivity. We have therefore performed simultaneous recordings in different regions of the primate dorsal visual pathway. Specifically, we recorded neural activity from the medial superior temporal (MST) area, and one of its main feedback targets, the middle temporal (MT) area. We estimated functional connectivity from correlations in the single-neuron spike trains and performed electrical microstimulation in MST to determine its causal influence on MT. Both methods revealed that inhibitory feedback occurred more commonly when the source and target neurons had very different stimulus preferences. At the same time, the strength of feedback suppression was greater for neurons with similar preferences. Excitatory feedback projections, in contrast, showed no consistent relationship with stimulus preferences. These results suggest that corticocortical feedback could play a role in shaping sensory responses according to behavioral or environmental context.

## Introduction

The functional organization of the visual system is often described in feedforward terms, as a cascade of processing stages that begins in the retina and continues through the thalamus to the primary visual cortex (V1) ^1-3^. V1 sends further projections to extrastriate cortical areas, which exhibit increasing selectivity for complex stimuli, while dependence on other stimulus properties recedes ^4,5^. The properties of these feedforward connections have been thoroughly studied, and indeed they now form the basis for a number of successful models of visual processing ^5-7^.

At the same time, anatomical studies have routinely found that feedforward inputs, despite their high synaptic weights, comprise a minority of the connections between visual areas. For instance, only 5% of synapses in the lateral geniculate nucleus (LGN) come from the retina, with 95% coming from the cortex and other brain structures ^8^.

Theoretical models have posited various roles for feedback computations ^9-12^. In many models, the strength and sign of feedback effects are related to the selectivity of the individual neurons in each area. For example, predictive coding models hypothesize feedback inhibition between neurons with similar preferences ^9^, while biased competition models hypothesize feedback excitation for similarly tuned neurons ^10^. Given the multiplicity of connections within the cortex, it can be difficult to distinguish between these possibilities ^11^.

Here we have examined the relationship between visual cortical stimulus preferences and feedback connections, using recordings from the middle temporal (MT) and medial superior temporal (MST) areas of non-human primate visual cortex. These areas encode visual motion defined by well-known stimulus parameters ^5,13-15^. By recording simultaneously from MT and MST, we were able to estimate functional connectivity in both the feedforward and feedback directions. We found that presumptive feedback connections from MST to MT were most often excitatory. However, the ensemble of these excitatory connections was unrelated to neuronal stimulus preferences. In contrast, inhibitory functional connections, while less frequently encountered, exhibited a clear preference for neurons with dissimilar stimulus tuning. Electrical microstimulation in area MST also tended to inhibit activity in area MT in a manner that depended on the relative stimulus preferences of the MT and MST sites and the strength of ongoing visual stimulation.

## Results

We performed simultaneous recordings from areas MT and MST in two rhesus macaque monkeys. The task involved simple fixation, and a large battery of visual motion stimuli (See Methods for details; ^5,15^) was used to characterize neuronal selectivity. We first present results that relate this selectivity to functional connectivity between pairs of neurons, as assessed with correlational methods ^16-18^. We then present the results of experiments involving causal manipulation of feedback, using electrical stimulation of area MST.

### Functional connectivity

We assessed the role of feedback connectivity in the dorsal visual pathway by recording simultaneously from areas MT and MST (Fig. 1a; blue and yellow arrows). These areas follow the classical cortical hierarchy, with MST receiving feedforward projections from MT and sending feedback projections to MT ^19^. To minimize stimulus-driven changes in connectivity, we first computed functional connectivity during periods of steady fixation on a blank screen. We then measured stimulus selectivity across space and feature domains on the same trials, using a wide variety of 100% coherent random-dot stimuli (Fig. 1b; see Methods for details).

**Figure 1:**
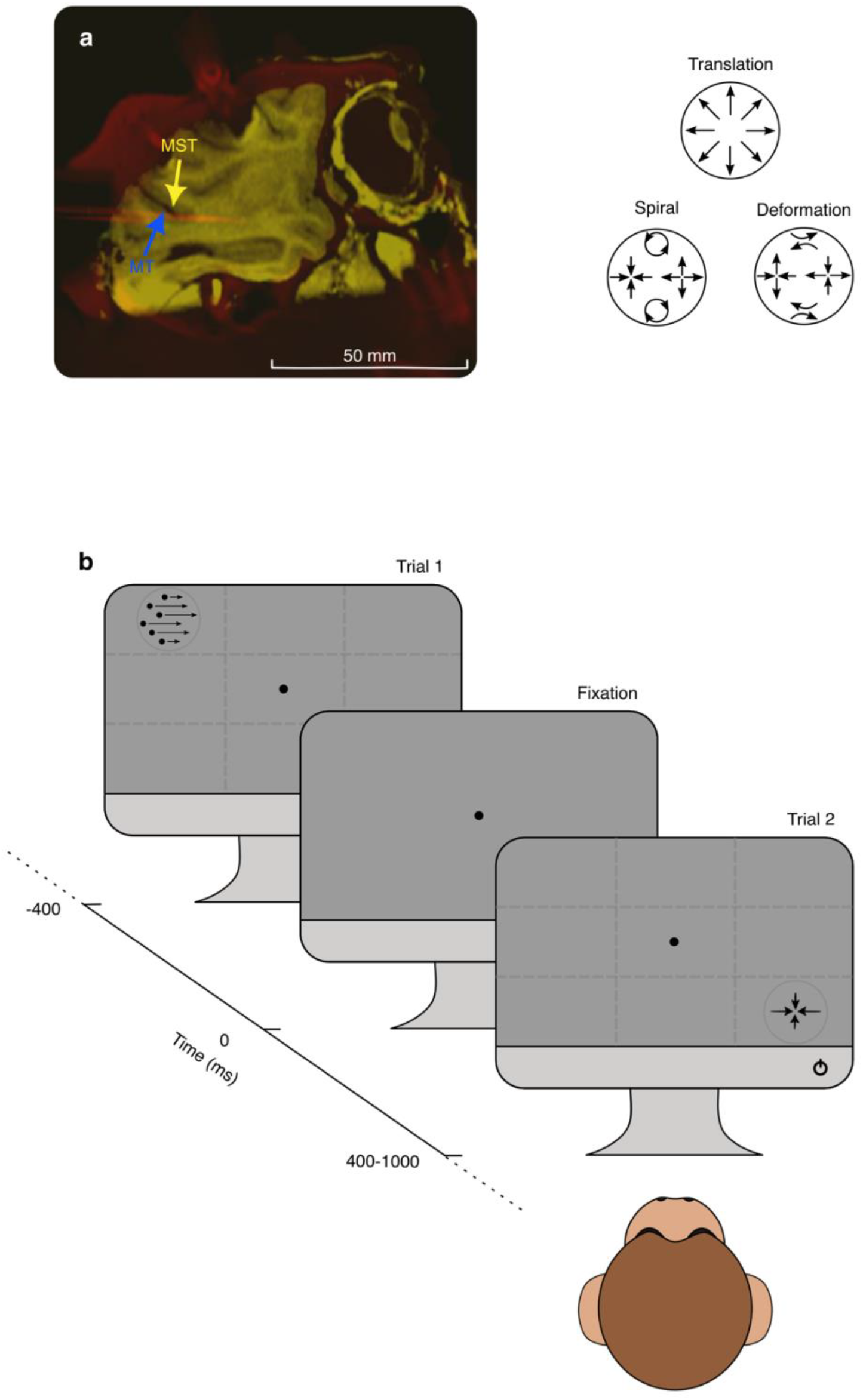
Electrophysiological recordings and experimental paradigm. **a**, Sagittal view of the MRI from one monkey. The orange lines indicate the tracks of the recording probe captured by a CT scan and registered with the anatomical MRI. The locations of MT and MST are indicated by blue and yellow arrows, respectively (adapted from ^77^). **b**, During each trial, the monkey fixated on a small dot positioned on the center of screen for a period of 400 ms. An RDK stimuli was then presented at one of the 9 possible locations on the screen for 400 ms to 1000 ms, depending on the animal’s capacity to work. The RDK stimuli consisted of translation, spirals, and deformation.

To identify pairs of neurons that exhibited significant functional connectivity, we first computed cross-correlations between the spike trains obtained from neurons in each pair of cortical areas. Cross-correlations capture how well the time series response from one neuron predicts the time series response of another neuron. As such, it provides a useful estimate of information flow from one cortical area to another ^16-18^. We then compared the stimulus selectivity of connected neurons, to infer possible roles for feedback and intermediate projections.

To compute cross-correlations, we first implemented a standard jitter correction procedure described elsewhere (see Methods for details; ^18,20^) to filter out the influence of slow temporal correlations. From the jitter corrected cross-correlations, we then extracted several relevant parameters, which are illustrated in Figure 2.

**Figure 2:**
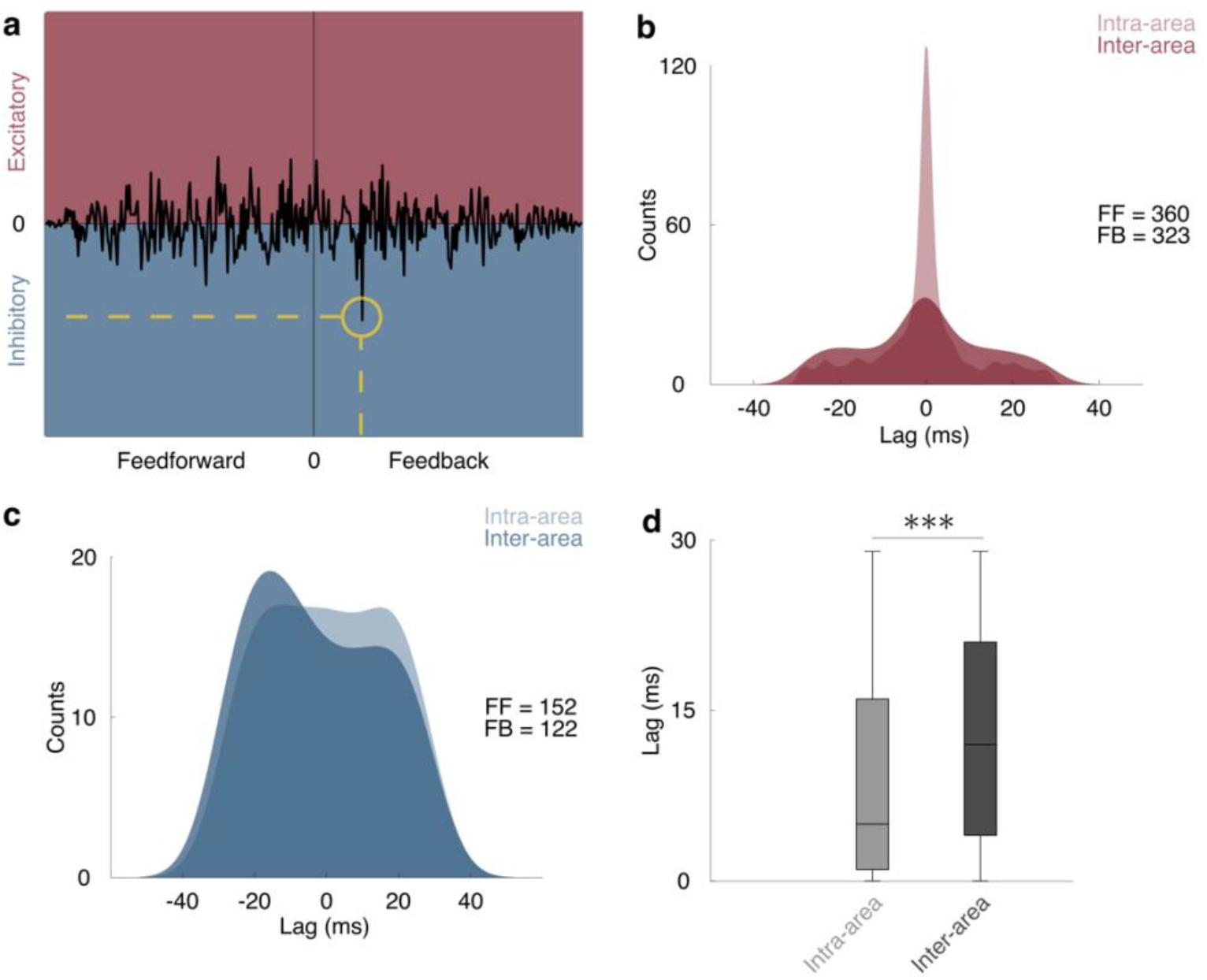
Cross-correlation as a measure of functional connectivity. **a**, Illustration of cross-correlation. Positive correlation peaks along the ordinate indicate excitation (red), and negative peaks indicate inhibition (blue). The location of the peak along the abscissa determines the direction of the connection. Peaks located to the left of the origin indicate feedforward connections, while peaks located to the right of origin indicate feedback connections. An example peak surrounded by the yellow circle represents an inhibitory feedback connection from area MST to MT. **b**, Comparison between intra-area (light red; MT-MT) and inter-area (dark red; MT-MST) significant excitatory functional connections. A WRS test did not reveal a significant difference between intra-area and inter-area excitatory connections (p = 0.96). **c**, Same as **b**, except for inhibitory connections (blue). A WRS test did not reveal a significant difference between the two categories (p = 0.96). **d**, Boxchart compares the distributions of lag for all connection types in intra-area (light gray) and inter-area (dark gray) pairs. A Two-sample t-test revealed a significantly lower mean latency for intra-area pairs compared to inter-area pairs (μ = 8.8 ± 0.01 ms vs. μ = 12.6 ± 0.01 ms; p =≪ 0.001). Whiskers represent lowest and highest lag values, and boxes represent upper and lower quartiles.

The lag of the jitter corrected correlation provides information about the potential direction of the connection: When an MT neuron consistently fires after an MST neuron (Fig. 2a, right side of the origin), for example, the functional connection is designated as feedback. Conversely, when the MST neuron spikes after the MT neuron, the connection is designated feedforward (Fig. 2a, left side of the origin). Additionally, from the sign of the correlation we can infer the type of connections, with positive correlations implying an excitatory connection (Fig. 2a, a global peak that appears in the red area) and negative correlations indicating an inhibitory connection (Fig. 2a, a global peak that appears in the blue area). Finally, the magnitude of the correlation lag can indicate if the source of the correlation is common input (zero lag), a monosynaptic connection (lags shorter than 2.5 ms or 4.5 ms), or a polysynaptic connection (lags longer than 2.5 ms or 4.5 ms) ^21-24^. We computed cross-correlograms (CCGs) for MT-MST pairs (*n* = 23,007) during the fixation periods before stimulus onset. We applied conservative criteria for identifying significant connections (see Methods), and as a result only 4.64% of the MT-MST CCGs were considered for further analysis.

We first examined excitatory functional connections. As shown in Figure 2b, these connections exhibited a mode at zero lag, indicating common input, for both intra-area (MT-MT; light red) and inter-area (MT-MST; dark red) pairs. Excluding pairs with zero lag (*n* = 109), we detected more excitatory feedforward (*n* = 360) than feedback (*n* = 323) connections.

Figure 2c shows the corresponding lag magnitude distributions for inhibitory functional connections for intra-area (light blue) and inter-area (dark blue) connections. These exhibited a wider range of correlation lags than the excitatory connections across both intra-area and interarea pairs. Excluding pairs with zero lag (*n* = 3), we again found that significant inhibitory feedforward connections (*n* = 152) were more common than their feedback counterparts (*n* = 122). Figure 2d shows that, as expected, the mean lag magnitude for intra-area MT pairs (light gray; 8.8 ± 0.01 ms) was significantly lower than for inter-area connections (dark gray; and 12.6 ± 0.01 ms; *p* ≪ 0.001; Two-sample t-test).

### Specificity of functional connections

We determined each neuron’s stimulus selectivity by presenting a large battery of motion stimuli that included 8 different directions of translation, 8 spiral optic flow stimuli (expansion, rotation and intermediates), and 8 deformation stimuli (combinations of expansion on one axis and contraction on the other). Each stimulus was shown at 9 different positions in the visual field (Figure 1b). Figure 3a illustrates a subset of the data for one example MT neuron (left) and one simultaneously recorded MST neuron (right). The tuning curves are represented as mosaic wheels, each of which corresponds to one spatial location (out of 9 possible stimulus locations) for the 8 directions of translation motion. Colors represent the firing rates for each motion direction, with red indicating high firing rates, white indicating firing near baseline levels, and blue indicating firing rates below baseline. The plot indicates that the MT neuron prefers motion upward and rightward and a stimulus position near the upper-left part of the visual field. The example MST neuron prefers leftward or upward motion, for stimuli in the left visual field. These neurons exhibited significant functional connectivity, with negative correlations in spiking activity and a positive correlation lag, suggestive of an inhibitory feedback connection. From data of this kind, we quantified the similarity in preferences for visual motion and stimulus position across the population of significantly connected neurons (see Methods).

**Figure 3:**
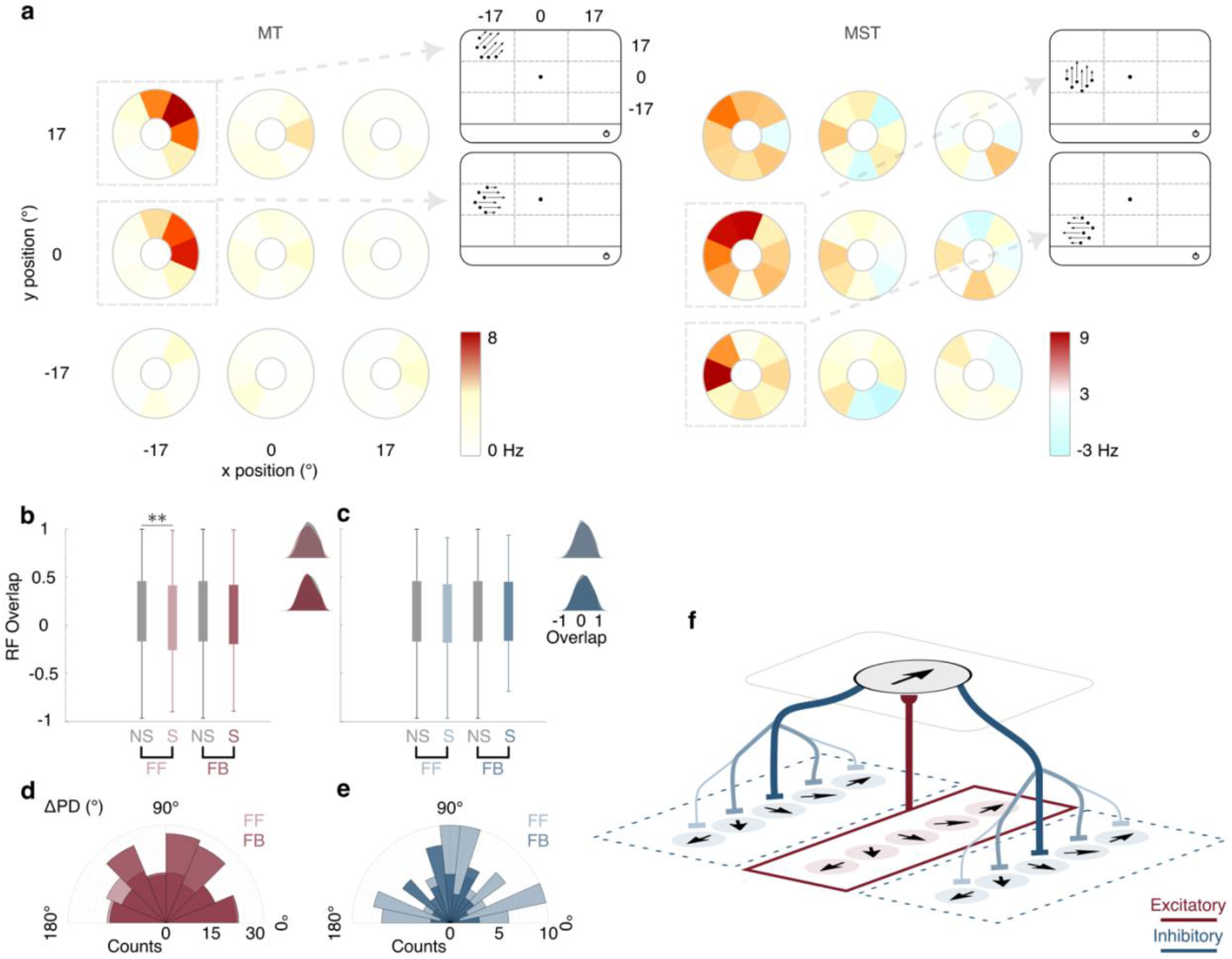
Spatial and feature tuning of functional feedback connections. **a**, Example tuning of a connected MT and MST neuron pair. Each motion type can appear in one of the nine possible stimulus positions (tuning wheels; shown with circle) on the screen. At each position, tuning wheels represent the neuron firing rate to eight motion directions; white represents baseline, red and blue correspond to highest and lowest firing rates compared to baseline. Inset shows the position and direction of a neuron’s tuning. Here, MT neuron prefers rightward and upward motion for stimuli positioned in the upper left, while MST neuron is tuned to leftward motion for stimuli positioned on the left of fixation. **b**, Boxchart compares RF overlap between non-significant (non-connected pairs; shown in gray) and significant (connected pairs; shown in red) excitatory connections in the feedforward (light red) and feedback (dark red) direction. A WRS test revealed significant RF overlap for excitatory feedforward connections compared to the non-connected pairs (p = 0.01; p = 0.17 for excitatory feedback connections). Whiskers represent the range of RF overlap. Zero indicates non-overlapping RFs and ±1 indicates highest degree of overlap (+1 yields a similar sign, -1 yields a reverse sign). Insets show the RF overlap distributions. **c**, Same as **b**, except for inhibitory feedforward (light blue) and feedback connections (dark blue). A WRS test did not reveal a significant difference between non-connected and connected pairs in either the feedforward (p = 0.25) or feedback (p = 0.48) directions. **d**, Polar-histogram shows distribution of the angle difference between connected excitatory feedforward (light red) and feedback (dark red) connections. A Rayleigh test did not reveal a significant tuning of the excitatory connections in either feedforward (p = 0.29) or feedback (p = 0.11) direction. **e**, Same as **d**, except distribution of the angle difference between connected inhibitory feedforward (light blue) and feedback (dark blue) connections are shown. A Rayleigh test revealed significant tuning of inhibitory feedback connections, forming a Von Mises distribution centered at 92° (p = 0.02), but not for inhibitory feedforward connections (p = 0.72). **f**, Summary of functional connection. Area MST (upper plane) receives excitatory feedforward input from MT neuron (lower plane) when RFs overlap (red box), while it sends inhibitory feedback connections (blue curved lines) to MT cells with orthogonal direction selectivity. Arrows indicate the cell’s preferred direction. Darker blue shades indicate more numerous connections.

We first considered the importance of spatial selectivity by comparing the extent of receptive field (RF) overlap between simultaneously recorded MT and MST neurons (see Methods). The results of this analysis are shown in Figure 3b and 3c. Pairs of neurons that exhibited significant excitatory feedforward connections had greater RF overlap than non-connected pairs (Fig. 3b; *p* = 0.01; *Wilcoxon rank sum* [*WRS*] *test*), but there was no significant difference between connected and non-connected pairs for excitatory feedback connections (*p* = 0.17; *WRS test*). Moreover, we did not detect significant RF overlap for inhibitory connections in either the feedforward or feedback directions (*p* = 0.25 *and p* = 0.48, *respectively*; *WRS test*). These results therefore suggest that feedforward excitatory connections are determined in part by RF overlap.

We next examined selectivity for motion direction of the neurons in each connected pair by measuring the absolute difference between their preferred directions (*ΔPD*), for both excitatory and inhibitory connections (see Methods for detailed procedure). We then compared these differences across the populations of feedforward and feedback connections. Figures 3d and 3e show the distribution of the difference in preferred directions for both excitatory (Fig. 3d; red polar-histogram) and inhibitory (Fig. 3e, blue polar-histogram) connections, across feedforward (light shades) and feedback (dark shades) connections.

For the population of excitatory connections (Fig. 3d), there was no significant bias toward neuronal pairs with similar or dissimilar motion preferences, in either the feedforward or feedback directions (*p* = 0.29 *and p* = 0.11, *respectively*; *Rayleigh test*). Inhibitory feedforward connections were also, as a group, untuned for motion preference similarity (Fig. 3e; *p* = 0.72; *Rayleigh test*). However, inhibitory feedback connections favored MT-MST pairs with a preferred direction difference near 90°, yielding a von Mises distribution that rejects the null hypothesis of a uniform distribution (Fig. 3e; mean 92°, *p* = 0.02; *Rayleigh test*, 95% confidence limits: 78.9° to 104.4°). Summary of these results are shown in Figure 3f; these confirm that inhibitory feedback connections from MST to MT are direction selective and mostly between cells with very different preferred directions (as in the example cell shown in Fig. 3a). Detailed summary statistics of these results are provided in Table 1 (RF_Overlap_) and Table 2 (*ΔPD*). A similar pattern of results was obtained when functional connectivity was computed during visual stimulation (Tables S1 and S2).

**Table 1.** Summary of statistics for RF overlap.

| **WRS test** | **MT → MST** | **MT ← MST** |
| --- | --- | --- |
| Excitatory | **p = 0.01** | p = 0.17 |
| Inhibitory | p = 0.25 | p = 0.48 |

**Table 2.** Summary of statistics for *ΔPD*.

| **Rayleigh test** | **MT → MST** | **MT ← MST** |
| --- | --- | --- |
| Excitatory | 91°<br>p = 0.29<br>CI = 81.3 to 100.2 | 85°<br>p = 0.11<br>CI = 78.4 to 93.5 |
| Inhibitory | 80°<br>p = 0.72<br>CI = 66.8 to 92.8 | **92°**<br>**p = 0.02**<br>**CI = 78.9 to 104.4** |

### Causal analysis of feedback projections

Thus far, our analysis suggests a common pattern of connectivity between individual neurons within the visual cortex, a key aspect of which is inhibitory feedback connectivity (Figure 3f). However, under natural conditions individual neurons are never activated in isolation, as even simple stimuli trigger responses in distributed networks of neurons. To examine feedback influences in this context, we presented various motion stimuli (Fig. 4a) and perturbed the resulting neuronal activity by electrically microstimulating in area MST (Fig. 4b). This allowed us to assess feedback influences through single-neuron recordings in MT (Fig. 4b). Microstimulation was delivered as brief pulses (5 pulses at 200 or 400 Hz), starting 150 ms after the onset of visual stimulation ^25^.

**Figure 4:**
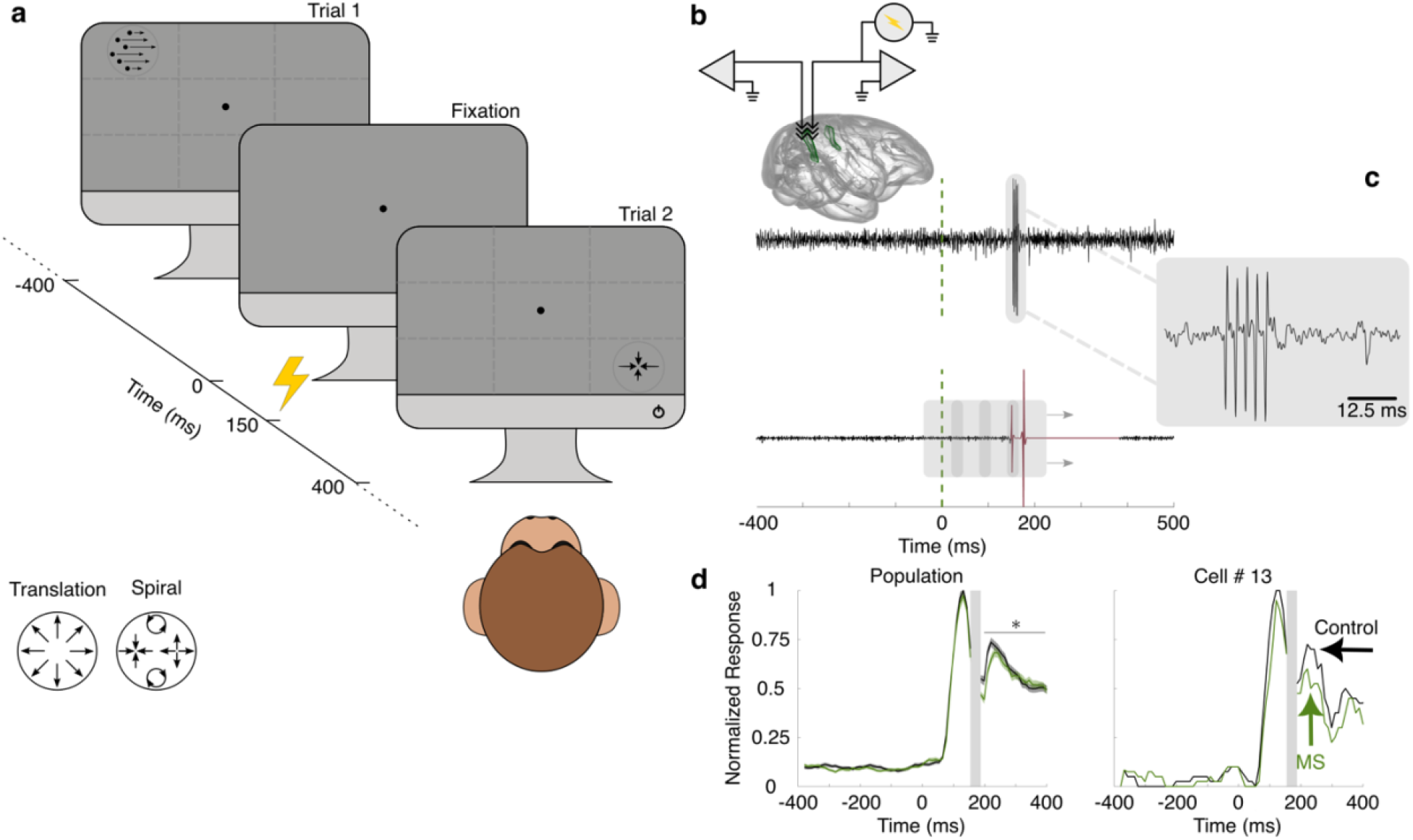
Causal connectivity experimental paradigm and artifact removal. **a**, Experimental setup is similar to **Fig.1b**, except for the application of electrical current 150 ms after stimulus onset (marked as zero on time axis) to causally perturb neuronal activity in area MST. Yellow flash indicates electrical microstimulation. **b**, Brief electrical currents were applied (5 pulses at 200 or 400 Hz) to area MST, while simultaneously recording were obtained from both areas MT and MST (green area) using two parallel micro-electrodes (triangles with arrows). A schematic of the brain is adapted from ^78,79^. **c**, Two types of artifacts detected in electrophysiological recordings. Type I artifact is shown with its specific characteristic including five large spikes locked to the microstimulation onset (upper panel). A snippet of broadband signal is shown in the gray box where electrical stimulation induced five artificial spikes and did not bleed through the remainder of the trial. Type II artifact shown in the lower panel resulted in a long-lasting pre-amplifier blackout (highlighted in red). Trials with Type II artifacts were detected using a 20 ms wide sliding window (gray boxes) and were excluded from the analysis pipeline. Dashed green line indicates visual stimulus onset. **d**, Left panel shows average MT neuronal response for control (black rate) and MS conditions (green rate). Shaded area surrounding each condition corresponds to a 95% confidence interval. Zero on the abscissa corresponds to stimulus onset, and the gray area reflects the microstimulation period (shown according to a 200 Hz stimulation), which was excluded from the analysis across both conditions. Microstimulation induces suppression across the MT population response, where the effect is statistically significant (p = 0.004; paired-sample t-test) for a period of 150 ms following microstimulation offset. Firing rate changes for an example MT cell are shown in the right panel.

As described in the Methods, microstimulation caused two types of artifacts on broadband signals. Type I artifacts reliably resulted in a series of large spikes time locked to the microstimulation onset on the broadband signal (Fig. 4c; top signal), while type II artifacts caused pre-amplifier saturation, resulting in a long-lasting blackout (Fig. 4c; bottom signal). Both types of artifacts have been detected by previous studies ^25,26^. We developed a simple yet effective procedure to detect corrupted trials using a sliding time window (Fig. 4c; gray moving boxes illustrate sliding detective windows; red trace shows pre-amplifier blackout). We then excluded those trials from the analysis pipeline. In total, we analyzed the visual responses of 84 single-units in area MT and 22 multi-unit clusters in area MST.

Figure 4d compares the average firing rate across stimuli, for an example MT neuron (right panel), for the control (black) and microstimulation (green) conditions. Consistent with previous work in extrastriate cortex ^25^, microstimulation caused a decrease of firing rates after stimulation offset. We observed a similar pattern of suppression induced by microstimulation when averaging the normalized firing rates of the entire MT population (Fig. 4d; left panel). The observed reduction in firing rate after microstimulation offset was significant for a period of 150 ms (6% decrease in magnitude; *p* = 0.004; paired-sample t-test).

### Specificity of feedback effects

Figure 5a shows the firing rate change in MT caused by microstimulation of MST as a function of the difference in preferred directions (*ΔPD*) for each MST-MT pair (*n* = 388). Overall, inhibitory effects (Fig. 5a, left; blue stars) of microstimulation are more common (65.46% or 254 pairs) than excitatory effects (Fig.5a, left; red stars) (box charts; Fig.5a, right; *p* ≪ 0.001; chi-squared test). As in the functional connectivity results (Figure 3), the inhibitory effect of microstimulation is more common for pairs with larger *ΔPD*s (Fig. 5a, right; *μ*_*I*_ = 104.4°; *p* = 0.005, *Rayleigh test*), while excitatory effects are distributed in a random fashion with respect to *ΔPD* (*μ*_*E*_ = 75.7°; *p* = 0.2, *Rayleigh test*). Interestingly, while inhibitory effects were more common for larger ΔPD, these effects were stronger for smaller ΔPD (blue line, Fig. 5a, left; *R*^2^ = 0.04; *p* = 0.002). As in the functional connectivity data (Figure 3), excitatory effects showed no consistent relationship with ΔPD (red line, Fig. 5a, left; *R*^2^ = 0.003; *p* = 0.83).

**Figure 5:**
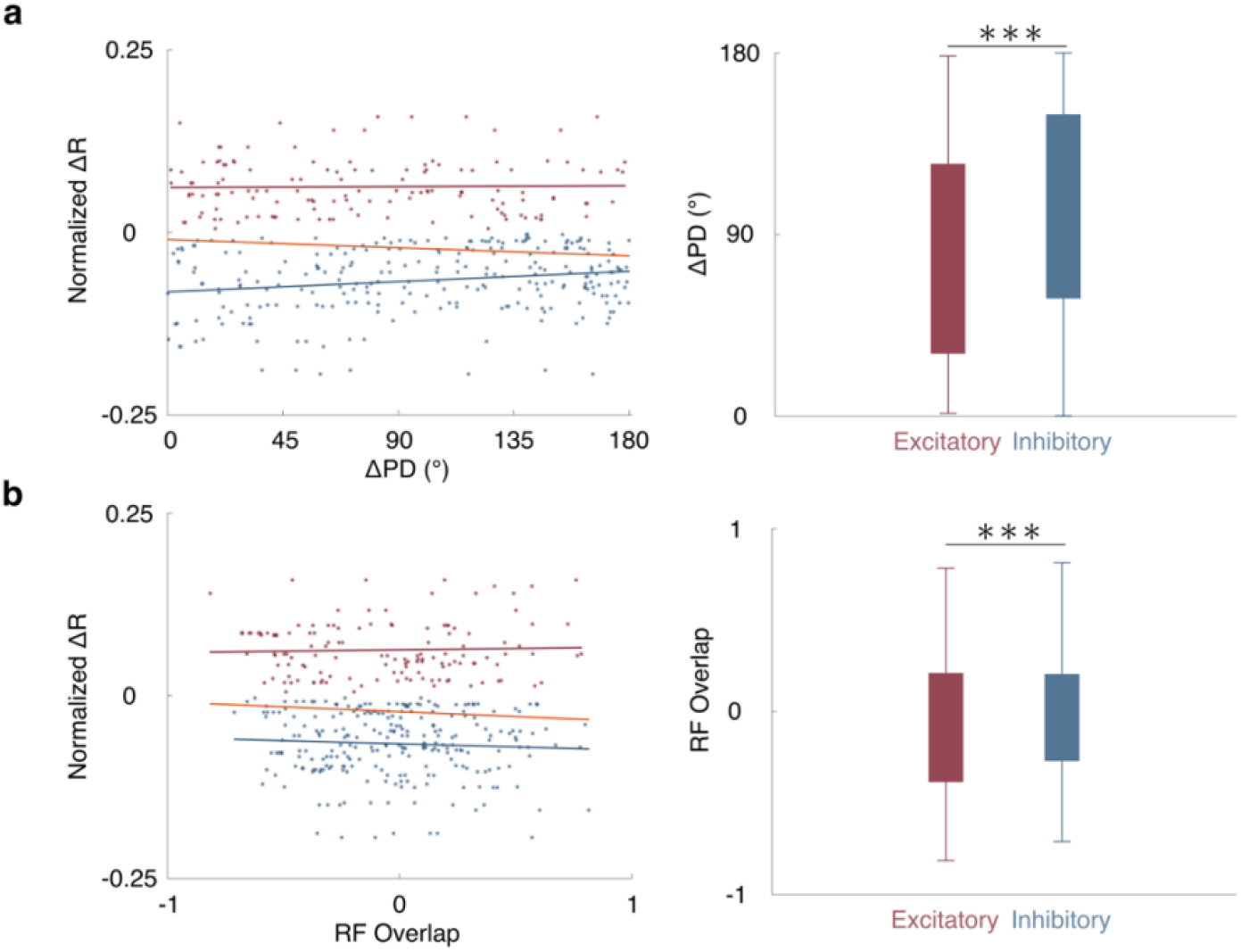
Specificity of feedback effects. **a**, MT firing rate changes after microstimulation as a function of MST preference. Left panel shows how individual MT neurons change their response as a function of ΔPD. Points below the abscissa indicate inhibition (blue stars), while those above the abscissa indicate an increase in response (red stars). Linear regression fit did not reveal a systematic influence from area MST (orange trend line; *R*^2^ = 0.008; *p* = 0.07). Red and blue trend lines represent a linear regression fit for excitatory (*R*^2^ = 0.003; *p* = 0.83) and inhibitory (*R*^2^ = 0.04; *p* = 0.002) effects. Boxcharts on the right panel compare MT neuronal response excitation (red bar) and inhibition (blue bar) across different ΔPD. A chi-squared test indicates that neuronal response inhibition is more likely than neuronal response excitation (p ≪ 0.001). Whisker represents the range of MST-MT preferred direction difference. **b**, same as **a**, except for RF overlap. Left panel shows individual MT neuronal response changes as a function of RF overlap between MST and MT. Linear regression fit did not reveal a systematic influence from area MST (orange trend line; *R*^2^ = 0.004; *p* = 0.2). Red and blue trend lines represent linear regression fit for excitatory (*R*^2^ = 0.001; *p* = 0.1) and inhibitory (*R*^2^ = 0.004; *p* = 0.3) effects. Boxcharts on the right panel is similar to boxchart in a, except it compares MT neuronal response changes across different RF overlaps. Whisker represents the range of MST-MT RF overlaps. A chi-squared test indicates that neuronal response inhibition is more common than neuronal response excitation (p ≪ 0.001).

We also examined the relationship between microstimulation effects and the relative RF positions of each MT and MST site (see Methods, Receptive field overlap section). Figure 5b shows the firing rate change in MT as a function of the RF overlap. Unlike the results for ΔPD, we did not observe systematic change on firing rates due to microstimulation across different RF overlaps for pairs of neurons that showed either excitation or inhibition (red line; *R*^2^ = 0.001; *p* = 0.1; blue line; *R*^2^ = 0.004; *p* = 0.3). These results suggest that inhibitory feedback influences are more common between sites with different stimulus preferences but stronger between cortical sites with similar stimulus preferences; in contrast, excitatory influences are relatively unspecific.

### Interaction of feedback perturbations with sensory-driven activity

We next considered the possibility that feedback effects depend on the strength of the visual stimulus, as has been suggested previously from single-neuron recordings and theoretical models ^27-32^. We first measured each MT neuron’s responses to our battery of motion stimuli in the absence of microstimulation and ranked the stimuli from weakest (lowest firing rate) to strongest (highest firing rate). For each stimulus rank, we then calculated the average firing rate difference (*ΔR* = *R*_*MS*_ − *R*_*Control*_) attributable to microstimulation. Overall, microstimulation resulted in a strong inhibitory effect that grew in proportion to the activation generated by the visual stimulus (Fig. 6a, left; linear regression fit; *R*^2^ = 0.67; *p* ≪ 0.001). For weaker visual stimulation, we observed an excitatory effect on neural responses. This transition from excitatory to inhibitory influences is similar to what has been seen previously across variations in stimulus contrast ^27-32^. We then sorted the MT neurons’ responses according to the ranking of the stimuli obtained at the MST site. When stimulus preferences were defined relative to the MST site, there was a significant but very small inhibitory influence of microstimulation for weaker visual stimulation (Fig. 6a, right; linear regression fit; *R*^2^ = 0.08; *p* ≪ 0.001).

**Figure 6:**
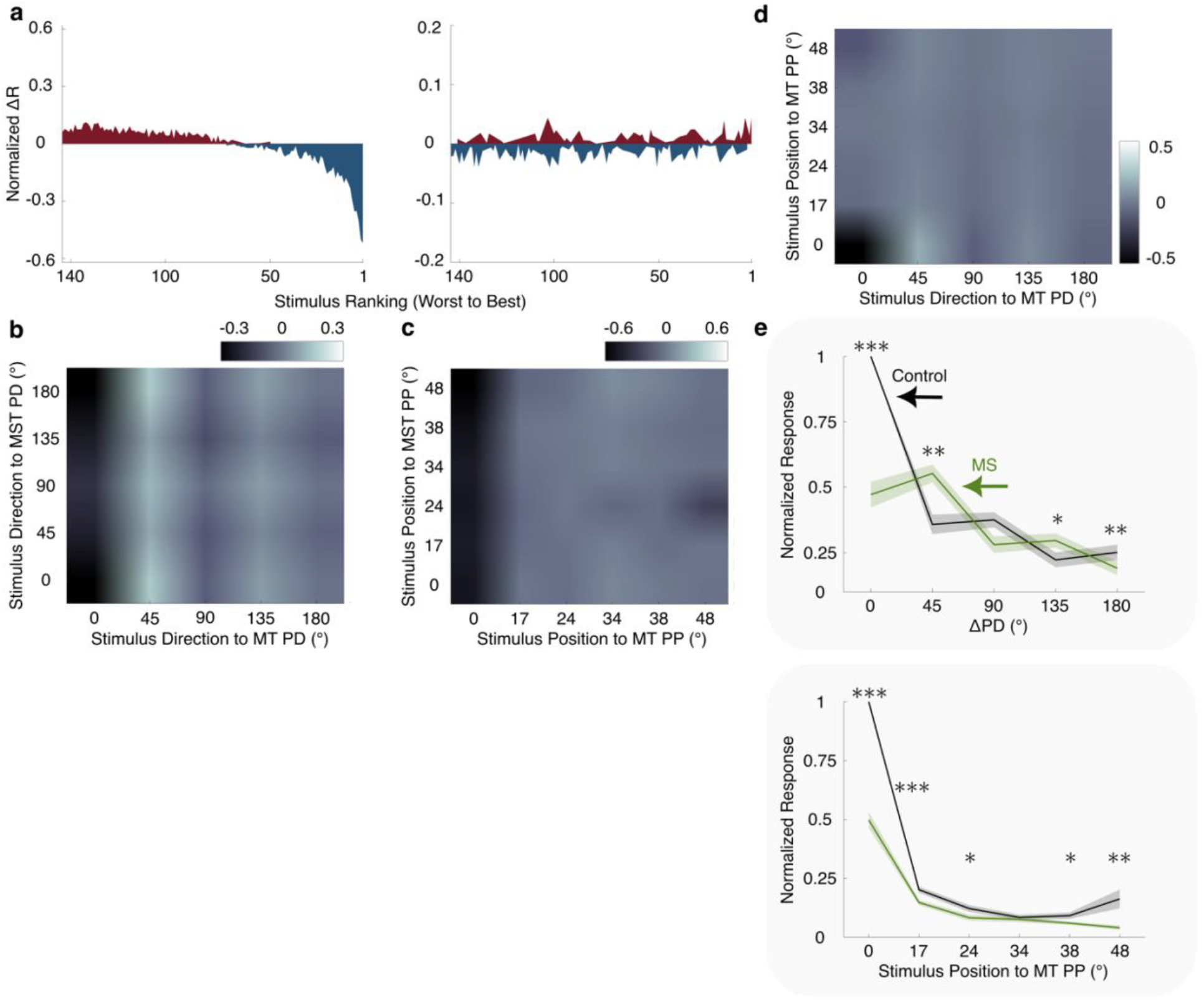
Modulatory effect of feedback connections. **a**, MST microstimulation effect on MT neural response to a battery of motion stimuli (total of 144 visual stimuli). Left panel: difference in MT neural response to different visual stimuli is ranked from weak (lowest response) to strong (highest response). The effect of microstimulation changes gradually from excitation (blue) for stimuli that elicited a weak response in the MT neuron to inhibition (blue) for those stimuli that elicited the strongest response in the MT neuron. Right panel: MT neural response to different visual stimuli is ranked based on MST preference. **b**, 2D matrix representation of MT ΔR in reference to MST PD (y-axis) and MT PD (x-axis). Colorbar represents the change in MT firing rate. **c**, similar to **b**, except for position selectivity. Stimulus position relative to MT preferred position is shown on the x-axis, while stimulus position relative to the MST preferred position is shown on the y-axis. **d**, similar to **b** and **c**, but for visual stimuli relative to MT only, when sorted in both direction (x-axis) and space domains (y-axis). **e**, MT neural response during control (black) and MS (green) to a wide range of visual stimuli at the best spatial location (top). A paired-sample t-test revealed significant effects of microstimulation for all ΔPDs’ except at 90^°^. Bottom panel, same as top panel but for MT neural response for the best stimuli at every possible spatial location. Shaded area surrounding each neuronal response corresponds to a 95% confidence interval.

To illustrate the overall pattern of excitatory and inhibitory influences, we constructed two-dimensional (2D) matrices of microstimulation effects on MT firing rates (*ΔR*, defined above). In these matrices one dimension captures the response relative to MT stimulus preferences, while the other dimension captures the responses relative to the preferences of the MST stimulation sites (see Methods for details). The results show that when area MT is probed with an optimal stimulus, microstimulation on average has an inhibitory effect that is largely independent of the preference of the MST sites (Fig. 6b; black colors). Instead, the effects are strongly dependent on the strength of the visual input relative to the selectivity of the MT population (Fig. 6b; first column on the x-axis). As in Fig. 6a, this inhibitory effect is replaced by excitation for the weakest stimuli (lighter colors). Figure 6c shows that similar results were obtained when stimuli were ranked according to the preferred positions (i.e. the RFs) of the MT or MST sites, while Figure 6d summarizes the effects of stimulus strength relative to MT preferences for both the direction and space domains.

To summarize the main conclusions of this section, Figure 6e (top panel) shows the average MT response for motion stimuli placed at the optimal spatial location for each site during control (black) and microstimulation (green) conditions. It reveals a significant reduction in firing rate for near-optimal visual stimuli (52.7% drop in firing rate at 0° deviation from PD; *p* ≪ *0*.*001*; *paired* − *sample t* − *test*). However, for weaker visual stimulation, there is a significant increase in firing rate for some stimuli (45°, *p* = 0.01; 135°, *p* = 0.04; 180°, *p* = 0.008; paired-sample t-test; <u>Table 3</u>).

**Table 3.** Paired-sample t-test summary for MT PD.

| **$\Delta PD$** | **0°** | **45°** | **90°** | **135°** | **180°** |
| --- | --- | --- | --- | --- | --- |
| Control vs. MS | **p << 0.001** | **p = 0.01** | p = 0.13 | **p = 0.04** | **p = 0.008** |

Figure 6e (bottom panel) shows the same analysis for the optimal motion stimulus placed at different spatial locations relative to the estimated preferred position of each MT cell. Again, there is a strong inhibitory effect of microstimulation that decreases with increasing values of ΔRF (see Table 4 for summary statistics). Our results at the population level therefore suggest that the influence of feedback connections interacts strongly with the instantaneous firing rate of the recipient cells, as determined by the strength of the visual input.

**Table 4.** Paired-sample t-test summary for MT PP.

| **$\Delta PP$** | **0°** | **17°** | **24°** | **34°** | **38°** | **48°** |
| --- | --- | --- | --- | --- | --- | --- |
| Control vs. MS | **p << 0.001** | **p << 0.001** | **p = 0.011** | p = 0.6 | **p = 0.04** | **p = 0.008** |

## Discussion

We have studied the relationship between feature selectivity and connectivity in the dorsal pathway of the primate visual cortex, using two different methods. The first is functional connectivity, a correlational method suitable for inferring the likelihood that any two individual neurons are connected via a relatively small number of intervening synapses ^16^. The results of our functional connectivity analysis suggest that tuned feedback connections tend to be inhibitory and between neurons with different stimulus preferences (Figure 3).

We also used electrical microstimulation to causally perturb activity in MST, while simultaneously monitoring the effects in MT. This revealed an effect that was also primarily inhibitory (Figure 5) and dependent on the strength of ongoing visual stimulation, with excitatory effects emerging for weak sensory input (Figure 6).

### Comparison with previous work

Historically, the anatomical evidence has yielded equivocal answers to the question of whether feedback connections were tuned to specific stimulus features ^33^. In non-human primates, most previous electrophysiological studies have used pharmacological and cooling interventions ^34-37^, which result in widespread inactivation of a cortical area. This makes it difficult to assess the functional specificity of individual feedback connections relative to stimulus selectivity. More recently Nurminen et al. ^38^ used optogenetic inactivation in anesthetized marmoset monkeys to selectively target individual V2 feedback inputs to V1. A consistent finding across all these studies has been an effect of feedback on RF size^34-38^, indicating a prominent role for feedback in regulating spatial summation. Similar effects have been reported in mice ^39-41^, with feedback tending to target cells with a high degree of spatial overlap ^39^, so as to sharpen the RF of the recipient cells while increasing surround suppression or contextual modulation ^34,41^.

Other studies have reported a relationship between stimulus selectivity and the retinotopic targets of feedback projections. In cats, feedback from V1 to the thalamus is determined in part by the orientation of the V1 cells and the retinotopic arrangement of targeted thalamic RFs ^42^. Similarly, corticocortical feedback to mouse V1 is often structured in retinotopic space according to selectivity for orientation or motion direction^39^. Marques et al. ^39^ further suggested that this kind of feedback is inhibitory, although they did not test this directly. Our work is consistent with this suggestion, with the caveat that the majority of the tuned inhibitory feedback connections in our data are likely polysynaptic connections (lags > 2.5 ms to 4.5 ms; ^21,22^). Thus, it is quite likely that some of the feedback influences we have described are mediated in part by lateral connections within MT.

Indeed, corticocortical feedback primarily targets excitatory neurons ^43^; the inhibitory influence of extrastriate feedback therefore occurs indirectly, via inhibitory interneurons ^44^. In V1, this inhibition is known to be regulated through local and long-range circuits in such a way that it depends on the strength of the feedforward input: The inhibition that is found for high stimulus contrasts turns to facilitation at low contrasts ^45-47^. A similar pattern of modulation is found in area MT ^28,29^. Our microstimulation results show similar effects (Figure 6), as perturbing feedback from MST to MT yields inhibitory effects when the feedforward influence of the visual stimulus is strong and excitatory effects when it is weak. Inhibitory surrounds likely facilitate figure-ground segmentation, while excitatory surrounds improve the encoding of stimuli under conditions of low visibility ^48^. The specific pattern of feedback influences we have found is also consistent with recent models of top-down attention to motion stimuli ^49^.

A recent macaque study has indicated that the relative contribution of feedforward and feedback connections changes during visual stimulation, with feedback connections being more dominant during the late phase of visual stimulation and periods of fixation ^50^. Similarly, our functional connectivity results show some evidence for changes in the relative impact of feedforward and feedback connections during strong visual stimulation (Tables 1S and 2S).

### Limitations of the current work

Both of our experimental approaches are limited in their ability to reveal the function of corticocortical feedback. For functional connectivity in particular, there is likely to be a bias toward detection of feedforward connections, which tend to be stronger than feedback ones. This might explain why we detected more feedforward than feedback pairs for both excitatory and inhibitory connections, despite the fact that, in primates, feedback connections outnumber their feedforward counterparts ^51^.

Similarly, microstimulation likely activated a patch of MST that included all neurons within a radius of a few hundred microns of the electrode tip ^52^. Such activation might have propagated within lateral circuits of MST. This kind of activity pattern is unlikely to occur with natural visual stimulation, and so our goal, as with previous work involving microstimulation ^53,54^, was to examine the effects of a relatively crude perturbation of ongoing activity.

Both the functional connectivity (Figure 3) and microstimulation (Figure 5) results revealed fewer connections between neurons with similar selectivity (small ΔRF or ΔPD), even though these connections appeared to be stronger than others (Figure 5). This might reflect the underlying reality of synaptic connections or a limitation of our indirect measurements. A more precise dissection of feedback influences is more feasible with other animal models. Rodents in particular are highly amenable to genetic and optical manipulations. However, the rodent visual system differs somewhat from that of the primate, with differences ranging from low visual acuity and lack of fovea ^55^, shallow visual hierarchy ^56^, and lack of cortical columns in V1 ^57^. Recent work with viral tracing methods in primates has revealed a clear organization of feedback projections between V2 and V1 ^58^.

### Theoretical implications

Feedback has been posited to play various roles in cortical computation. For the specific case of MST-MT feedback, Mineault et al. ^5^, showed that inhibitory feedback influences of the kind we have demonstrated empirically could subserve the computational function of sharpening selectivity to complex stimuli, while maintaining invariance to irrelevant features. Additionally, tuned inhibitory feedback is an important component of a number of theoretical models ^9,59,60^. In these models, feedback carries predictions about the expected activity of lower-level neurons. By inhibiting this activity, only the “prediction errors”, which are unexpected aspects of the sensory input, are propagated through the cortical hierarchy. This process is thought to lead naturally to observed phenomena such as attentional and contextual modulation of sensory responses ^9^.

More generally, biological neural networks respond to strong sensory inputs with strong excitation that is often followed by adaptation, usually in the form of an inhibitory response that maintains firing rates within a relatively narrow range ^61-64^. This kind of response regulation can be performed by standard models, such as normalization ^62^, but these models do not account for the kinds of increased neural responses that can occur in response to microstimulation (Figure 6). Instead, the kind of feedback we have found – inhibitory during strong ongoing activation and excitatory during weak activation – could play a role in homeostasis regulation, the universal process of maintaining physiological stability in the face of fluctuating environmental conditions. This function can be accommodated within the predictive coding framework ^9,65^.

Our results do not reveal the precise circuitry of feedback modulation. While we have shown that the ultimate influence of feedback depends on neural selectivity, we cannot say whether this selectivity is instantiated directly via the underlying anatomical connections, or determined by lateral connections within MT. These possibilities are notoriously difficult to distinguish with extracellular recordings ^11^. In the limit, it could be that feedback is entirely untuned, but it activates a network of tuned recurrent connections within the recipient area ^44^. Given that many of the influences we have detected are likely to be polysynaptic ^21,22^, it would be useful to examine the properties of models such as the supralinear stabilized network ^31,66,67^, which rely heavily on recurrent connectivity and have the explicit goal of maintaining cortical stability.

## Methods

### Electrophysiological Recordings

Two adult rhesus macaque monkeys (Macaca mulatta; 7 kg female-monkey Y and 18 kg male-monkey S) were used for electrophysiological recordings. Eye movements were sampled and monitored during recording sessions at a sampling rate of 1000 Hz using an EyeLink 1000 infrared eye tracking system (SR Research). Each monkey’s head was stabilized during the experiments using an MRI-compatible titanium head post that was implanted on the skull. All procedures abided by regulations set by the Canadian Council on Animal Care and were approved by the institutional Animal Care Committee of the Montreal Neurological Institute.

Broadband signals were recorded using a data acquisition system (Plexon Multichannel Acquisition Processor System). For MT neurons, single-unit (SU) spike sorting was performed by thresholding and filtering the wideband neural signal online, then clusters of spikes were assigned to single units using an automated template-matching algorithm (Plexon MAP System). This procedure was followed by manual spike sorting to obtain SU activity using a combination of automated template matching, clustering based on the principal components, absence of absolute refractory period (1 ms) violations (Plexon Offline Sorter), and visual inspection of waveforms.

### Functional connectivity

We conducted a functional connectivity experiment that involved simultaneous recordings in the superior temporal sulcus (STS), where area MT is located in the posterior side and MST is located in the anterior side ^13,14,68^. During each recording session, we approached the two areas posteriorly by placing a 32-channel linear microelectrode arrays (Plexon V-Probes with 0.15 mm inter-channel spacing and impedance range between 1-2MΩ) so that the more superficial 16 channels (spanning ∼2.4mm) were located within area MT, while the deeper 16 channels reached area MST. Both regions were identified based on anatomical MRIs, brain atlas, depth from cortical surface, transitions between white and gray matter, and a clustering algorithm explained in the next section (see Fig. 1a for traces of a typical recording path).

### Causal connectivity

For our causal connectivity experiment, we performed electrophysiological recording and electrical microstimulation in monkey Y using two 32-channel linear microelectrodes separated by approximately 1 mm gap from each other. Each microelectrode was positioned to cover both MT and MST (see Fig. 4b for the arrangement of recording probes). We drove the microstimulating electrode deeper to ensure a larger area of MST was covered. The areas were then identified as described in the functional connectivity experiment (see Fig. 1a for traces of recording path).

### Verification of recording sites

We confirmed electrode positions using computed tomography (CT) imaging. Stainless-steel guide tubes were inserted into the recording chamber, and 250 µm tungsten electrodes were placed 5 mm above the typical recording locations. We then acquired a single volume with 300 µm isotropic voxels using a Vimago CT scanner (80 kVp). The CT imaging data were registered with the anatomical MRI scans to precisely visualize the electrode locations (Fig. 1a).

### Visual Stimuli

An LCD screen with a 75 Hz refresh rate and a resolution of 1280 x 800 pixels was used to display motion stimuli spanning a viewing area of 60° horizontally and 40° vertically. For the functional connectivity experiment (Fig. 1b), the animals maintained their gaze at a fixation point in the center of the screen for a period of 400 ms. The visual stimulus was then presented for another 400 ms to 1000 ms, while the animals kept fixating at the center point. For the causal connectivity experiment (Fig. 4a), the stimulus presentation period was limited to 400 ms. Gaze was required to remain within 1.5° of the fixation point for the animal to receive a liquid reward. Data from trials in which gaze was not maintained in the fixation window were discarded.

The direction tuning and receptive field locations of single cells were assessed using 100% coherently moving white-dot stimuli on a black background. The dot density was 2 dots/deg.^2^, where each dot had a lifetime of 10 frames and a diameter of 0.1°. On each trial, a single patch of dots was displayed, centered at a location selected pseudo-randomly from a 3 by 3 grid of possible stimulus positions (Fig. 1b). The fixation point was located at the center of the 3 by 3 grid, as well as the center of the screen. The patch radius ranged from 6° to 15° based on initial hand mapping for the pool of cells found in each day. The separation between grid positions ranged from 10° to 17° with identical dimensions for both the vertical and horizontal axes. However, for the causal connectivity experiment, patch radius and separation between grid positions were 15° and 17°, respectively.

Directional tuning was identified for the 3 different motion types defined in the equations below: translation (Eq. 1), spirals (Eq. 2), and deformation (Eq. 3).

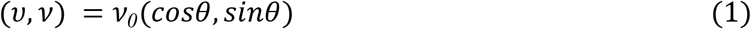

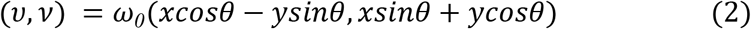

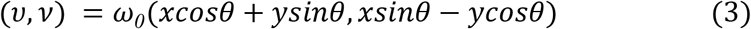

These types were chosen because they span the space of first-order optic flow ^69^. In each of these equations, (*u, v*) represents the instantaneous velocity of a dot at position (*x, y*). For each category of motion, we presented 8 directions, represented by *θ* in the equations below, sampled at 45° intervals. For translation motion, dots moved in one direction, and the speed (v_0_) ranged from 8 °/s to 32 °/s, though most recordings were performed at 16 °/s. For spiral stimuli, the 8 directions corresponded to expansion, contraction, rotation, and their intermediates. The speed (w_0_) was set to a value between 3 Hz and 5 Hz. For deformation, motion involved expansion on one axis and contraction on the perpendicular axis, with the axis varying across motion direction *θ*. The speed of motion for deformation was always the same as the speed of the spiral stimuli, which was selected based on an initial, qualitative screening of neuronal responses.

For the functional connectivity experiments, the combinations of all directions, motion types, and stimulus locations resulted in 216 stimuli that were randomly interleaved and displayed 4 or 5 times each. For the causal connectivity experiment, we used only the translation and spiral stimuli, for a total of 144 stimuli, each with 8 repetitions. We applied microstimulation for half of the repetitions, chosen pseudo-randomly between no-microstimulation and microstimulation conditions. Microstimulation was applied 150 ms after visual stimulus onset (Fig. 4a).

### Microstimulation Procedure

We causally manipulated neural activity using an electrical microstimulation device with a stim-switch controller and stim-switch head-stage (BlackRock MicroSystems). The microstimulation system was triggered by an activation pulse (5V TTL) from our recording system to the stimulator, which also served as a timing pulse to ensure accurate synchronization with the recording system. A train of 5 negative-first biphasic pulses (200 µs per phase) was applied to modulate the activity at the microstimulation site at a frequency of 276.9 ± 101.3 Hz and amplitude of 56.1 ± 9.6 µA (similar to ^25^). The mid-level current amplitudes chosen in our experiment likely activated a restricted set of cortical columns in area MST ^52^. Additionally, to ensure perturbation of feedback connections from MST to MT we triggered the micro-current train 150 ms after visual stimulus onset ^25^. In every recording session, we changed the microstimulation channel to prevent impedance degradation of the contact site. Finally, to designate an MST cell for microstimulation, we ensured the visibility of strong neural signals prior to stimulation.

### Data Analysis

All analyses were performed using built-in and custom-written code in MATLAB R2022b.

### Functional connectivity analysis

Firing rates were computed for each neuron by taking the mean spike rate averaged across stimulus repetitions over the period from 50 ms after stimulus onset to the end of stimulus period. Baseline firing rates were computed as the mean firing rate during the entire fixation period (400 ms), then averaged across all trials in a single task. Only visually responsive units, whose firing rate increased at least 2 standard deviations above the mean of the spontaneous firing during visual stimulation, were analyzed.

### MT to MST Transition

We used a clustering-based approach to determine the transition from MT to MST, based on the assumption that there is a brief range of low visual activity coinciding with the electrode’s transition from one side of the superior temporal sulcus to the other side. Additionally, we categorized MT and MST cells based on the spatial distribution of channels of significant visual activity (whose firing rate during visual stimulation increased at least 2 standard deviations above the mean of the spontaneous firing). The model assumes that MT cells are spread over the more superficial end of the electrode and MST cells are spread around the deeper end, during a posterior approach. A boundary *B* is determined by minimizing the sum of pairwise distances *d*_*p*_ between significantly responsive cells (represented by channel number *ch* in the equation below) within each cluster for the 2 centroids as follow:

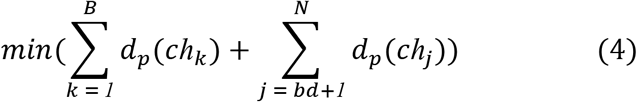

where *N* is the total number of channels in each recording.

### Cross correlation analysis

To estimate functional connectivity, we calculated the normalized spike train cross-correlogram between all pairwise combinations of neurons recorded in each session ^16,18,70^. For a given trial, we first calculated the cross-correlogram between two neurons as:

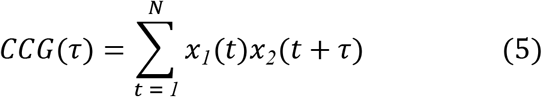

where *x*_*1*_(*t*) and *x*_*2*_(*t*) are the binary spike trains of the two neurons, sampled at 100 µs resolution, *N* is the number of bins in the trial, and *τ* is the time lag. The resulting cross correlations were then smoothed with a 5 ms sliding window Gaussian filter (represented by the function *G* in equation 6), and then averaged over all trials. Because spike trains carry stimulus locked information that are correlated across all neurons, we implemented the jitter correction procedure proposed in Smith and Kohn ^18^. The procedure is explained in full detail elsewhere ^18,20,71^, but briefly, the spikes were shuffled across trials within a jitter window of 50 ms, maintaining their original firing rate (number of spikes in each bin). The process was then repeated 1000 times for each pair of spike trains and a correction factor *CF* was computed by averaging the cross correlations calculated after jittering. By subtracting the correction factor *CF* from the cross correlation calculated without jittering, we obtain a jitter corrected *CCG*. The jitter corrected *CCG*s were then normalized by the geometric mean spike rate 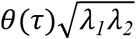 where *θ*(τ) is a triangular function that accounts for the extent of overlap between the spike trains at each time lag, and *λ*_1_ and *λ*_2_ are the mean firing rates for the pair of neurons ^17,18^. The corrected CCG is given by:

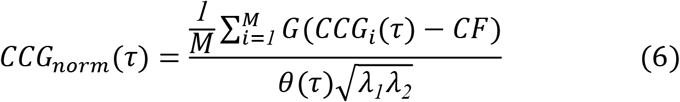

where *M* is the number of trials in the session.

To filter out the *CCG*s that did not represent significant connectivity between pairs of neurons, we only considered values of *CCG*_*norm*_(τ) where the absolute peak was at least equivalent to 5 times the standard deviation of the flanks. The flanks were defined as the tails of the cross correlations ranging from ±75 to ±150 ms, considering that the maximum lag at which the *CCG*(*τ*)s computed was 150 ms. We further excluded the *CCG*_*norm*_(*τ*)s where the absolute peaks occurred at or beyond 30 ms of lag, as studies showed that feedback signal affects neural activity in the downstream area on the early part of the response ^36,72^.

To ensure that the correlations we detected did not arise by noise or programming errors, we applied our analysis pipeline to pairs of spike trains artificially generated using a Poisson process. The firing rates of the artificial spike trains were selected randomly from a pool of firing rates obtained from the real data. The percentage of correlations that passed our criteria was 0.47%, suggesting the majority of connections found in our data are not due to the statistics of spiking or spurious correlations.

### Causal connectivity analysis

For both SU and MU activities during control and MS conditions, firing rates were computed for each neuron by taking the mean spike rate averaged across stimulus repetitions over the period from 50 ms after stimulus onset to the end of stimulus period excluding 15.38 ± 3.8 ms (mean ± SD) period corresponding to microstimulation period. Baseline firing rates were computed as the mean firing rate during the entire fixation period (400 ms), then averaged across all trials in a single recording session.

We excluded neurons whose firing rate during stimulus presentation was not significantly higher compared to spontaneous firing rate (*p* < *0*.*05*; paired-sample t-test). However, since electrical current propagates through cortical structure at the microstimulation site and activates a pool of neurons around the stimulation channel, we included MST cells that were located ± 400 µm away from the cell that received microstimulation ^52^.

### Signal Acquisition and Artifact Removal

Electrical microstimulation caused two types of artifacts on the raw voltage traces (Fig. 4c). Type I artifacts reliably resulted in 5 large spikes locked to the microstimulation onset (Fig. 4c; top panel), lasting for 15.38 ± 3.8 ms (mean ± SD), while Type II artifacts caused pre-amplifier saturation (amplitude ≪ 10^−19^) and resulted in a long-lasting blackout (Fig. 4c; red signal in bottom panel). In order to detect and remove these artifacts we first started by subtracting the DC offset from the raw voltage traces, then applying a Butterworth band-pass filter (300 - 3000 Hz; ^73^) to produce multi-unit (MU) activity signals for MST cells. MU signals represent the average spiking activity of neurons close to the recording channel on the microelectrode. Prior to removing artifacts, MU signals were segmented into trials. We detected the Type II artifacts from each trial by searching for blackout periods that lasted for a minimum of 20 ms and those trials were then excluded from analysis. Moreover, MU channels with multiple corrupted conditions were also excluded from analysis. Finally, MU spikes were obtained from waveforms that exceeded a certain threshold amplitude, defined as 3 times the standard deviation of the background noise estimated from the bandpass-filtered signal ^73^. To control for Type I artifacts, the corresponding microstimulation period was excluded from the spike train for both control and microstimulation trials.

### Receptive field overlap

To evaluate the spatial extent of the RF overlap between pair of neurons from two different areas (MT-MST), we measured the correlation coefficient between neuron 1 and neuron 2 by first normalizing each RF and then calculating the dot product of their average firing rates in response to all visual stimuli as a function of stimulus position defined below ^74,75^:

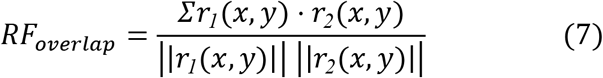

Here *r*_*1*_ *and r*_*2*_ are the mean firing rate of each neuron in a given pair, *x and y* correspond to stimulus coordinates, and |❘. ❘| denotes the norm of the response vector. The value of the *RF*_*overlap*_ ranges between -1 (identical RF with opposite sign) to +1 (identical RF with similar sign).

### Preferred direction difference

For each neuron, we formed three tuning curves, one for translation motion, one for spiral motion, and one for deformation motion. We only considered responses to stimuli presented at the position that elicited the maximum firing rate. For each pair of connected neurons, we then calculated the difference in preferred directions from one of the tuning curves (translation, spirals or deformation). We chose the motion type based on the preference of the neuron that lagged behind in the cross correlation. If the computed lag of maximum correlation reveals that neuron 1 follows neuron 2 in response, we use the preferred motion type for neuron 1 as a reference to compute directional selectivity. Neuron 1 in that case represents the termination of the assumed anatomical connection (approximated from the functional connection).

For this metric, we excluded pairs of neurons in which one or both were not direction selective. We defined direction selectivity based on a standard selectivity index:

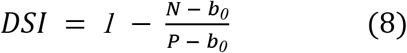

where *P* and *N* are the mean firing rates in the preferred direction and null directions, respectively, and *b*_0_ is the mean spontaneous (“baseline”) firing rate. To ensure meaningful measures of preferred direction, we required both neurons in the pair to have DSI > 0.7. We then estimated the preferred direction for each neuron from the circular mean 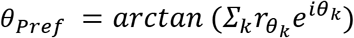, where 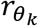 is the firing rate at angle *θ*_*k*_. The difference in preferred angle was then calculated by obtaining the circular difference between the preferred directions for the two neurons.

### Statistical analysis

All statistical analyses were performed using built-in MATLAB R2022b functions and/or custom scripts. We used the parametric paired-sample t-test to compare neural responses in two different conditions. Parametric two-sample t-tests were used to compare mean latency of intra-area and inter-area connections. The non-parametric Wilcoxon rank-sum test were used for RF overlap distributions and time lag distributions. We performed the Rayleigh test to identify Von Mises distributions for circular variables using the CircStat toolbox for MATLAB ^76^. Significance was defined as *p* ≤ *0*.*05*; all tests were two-tailed.

## Data availability

The data will be provided upon reasonable request to the corresponding author.

## Acknowledgements

We thank Julie Coursol for outstanding animal care and Melanie Segado for help with the summary figures and insightful comments on the paper. This work was supported by a grant from the CIHR (PJT178071).

## Author contributions

Y.K., N.N., and C.C.P designed the analyses. Y.K. and N.N. collected electrophysiological data.

Y.K. performed causal connectivity analyses. N.N. performed functional connectivity analyses.

Y.K. contributed to functional connectivity analyses. Y.K., N.N. and C.C.P wrote the manuscript.

C.C.P. supervised all aspects of project. All authors discussed the results and commented on the manuscript.

## Competing interests

The authors declare no competing financial interests.

## Supplementary information

**Table 1S.** Summary of statistics for RF overlap during visual stimulation.

| **WRS test** | **MT → MST** | **MT ← MST** |
| --- | --- | --- |
| Excitatory | p = 0.44 | **p << 0.001** |
| Inhibitory | p = 0.77 | p = 0.19 |

**Table S2.** Summary of statistics for *ΔPD* during visual stimulation.

| **Rayleigh test** | **MT → MST** | **MT ← MST** |
| --- | --- | --- |
| Excitatory | 77°<br>p = 0.9<br>CI = 66.6° to 86.9° | 90°<br>p = 0.46<br>CI = 79.6° to 100° |
| Inhibitory | **98°**<br>**p = 0.053**<br>**CI = 80.5° to 116.2°** | 70°<br>p = 0.25<br>CI = 50.1° to 90.4° |

## Notes

### Competing Interest Statement

The authors have declared no competing interest.

### Summary of Updates

Minor text edits. Section "Comparison with previous work" in discussion updated for clarification. Figure 5b has been updated.

